# Revealing the abiotic and biotic drivers of past and future local adaptation in a coastal dogwhelk

**DOI:** 10.64898/2026.08.21.746321

**Authors:** Emily K. Longman, Eric Sanford, Joaquin C.B. Nunez, Melissa H. Pespeni

## Abstract

Predicting whether populations can persist under rapid environmental change requires identifying the ecological drivers of local adaptation, uncovering their genetic bases, and understanding how adaptive variation will respond to future selection. Here, we combine landscape genomics, environmental data, and evolutionary simulations to identify the selective forces shaping adaptation across 1,500 km of the west coast of North America in the low-dispersing coastal dogwhelk *Nucella canaliculata*, determine their genomic bases, and forecast future evolutionary responses. We found strong associations of genome-wide variation with both abiotic (i.e., mean pH) and biotic variation (i.e., cross-sectional shell thickness of the mussel prey species, *Mytilus californianus*). These patterns are underlain by two large-effect loci, including a biomineralization gene associated with pH tolerance and a locus near a thiamine transporter associated with prey shell thickness. Genomic offset analyses and population genetic simulations further predict that ongoing ocean acidification will disrupt existing adaptive patterns and generate maladaptation in high latitude populations, with evolutionary outcomes strongly influenced by gene flow, which determines the rate at which adaptive alleles spread across the species range. Together, these findings reveal how biotic and abiotic selective pressures shape adaptive genomic variation and provide a framework for forecasting evolutionary responses to future global change.

## Introduction

Ecological conditions vary across landscapes, generating heterogeneous selective pressures that shape adaptive divergence of phenotypic traits and allele frequencies (1–4). However, it remains a challenge to identify the selective mechanisms and genomic architecture underlying patterns of local adaptation due to the complexities of natural ecosystems, historical demography, and variation in organismal life history (5). Further, insights into the mechanisms shaping contemporary adaptive variation are increasingly urgent, as climate change is modifying environmental conditions, reshaping species ranges and community assemblages, and altering patterns of biotic selection, yet the capacity of species to respond adaptively remains uncertain (6–9). Here, we study the genomic bases of local adaptation in a non-model coastal species to advance our understanding of the evolutionary processes driving local adaptation and to predict the vulnerability of populations to global climate change.

Rocky intertidal ecosystems have strong environmental gradients and are one of the most extensively studied marine habitats (10). The large body of work testing ecological theories in this coastal ecosystem (11–13) affords access to a deep understanding of species interactions and large-scale ecological data. This rich ecological foundation makes rocky intertidal habitats ideal natural laboratories for connecting ecological drivers of selection to genomic variation and for forecasting the fate of local adaptation under future environmental change. In this study, we leverage the biology of the channeled dogwhelk, *Nucella canaliculata*, as a system to identify the ecological drivers and genetic bases of local adaptation across 1,500 km of the west coast of North America. This dogwhelk lives on wave-exposed rocky headlands along the northeast Pacific and has low dispersal potential — females lay clusters of benthic egg capsules, and crawl-away juveniles settle in the nearby area (**Fig. 1A**; [14]). Recent work has indicated a high level of population structure (15) and locally adapted patterns of phenotypic variation, reflecting adaptation to both abiotic and biotic conditions (16–20). For example, *N. canaliculata* populations vary geographically in predatory drilling ability (14, 16–18), a slow and energetically costly chemo-mechanical process that has implications for mussel bed community dynamics (21–23). Variation in this phenotypic trait has been associated with the abundance of preferred prey – the blue mussel *Mytilus trossulus* and the acorn barnacle *Balanus glandula* (17) – and the shell traits of the most abundant mussel prey species, *M. californianus* (18).

**Figure 1.**
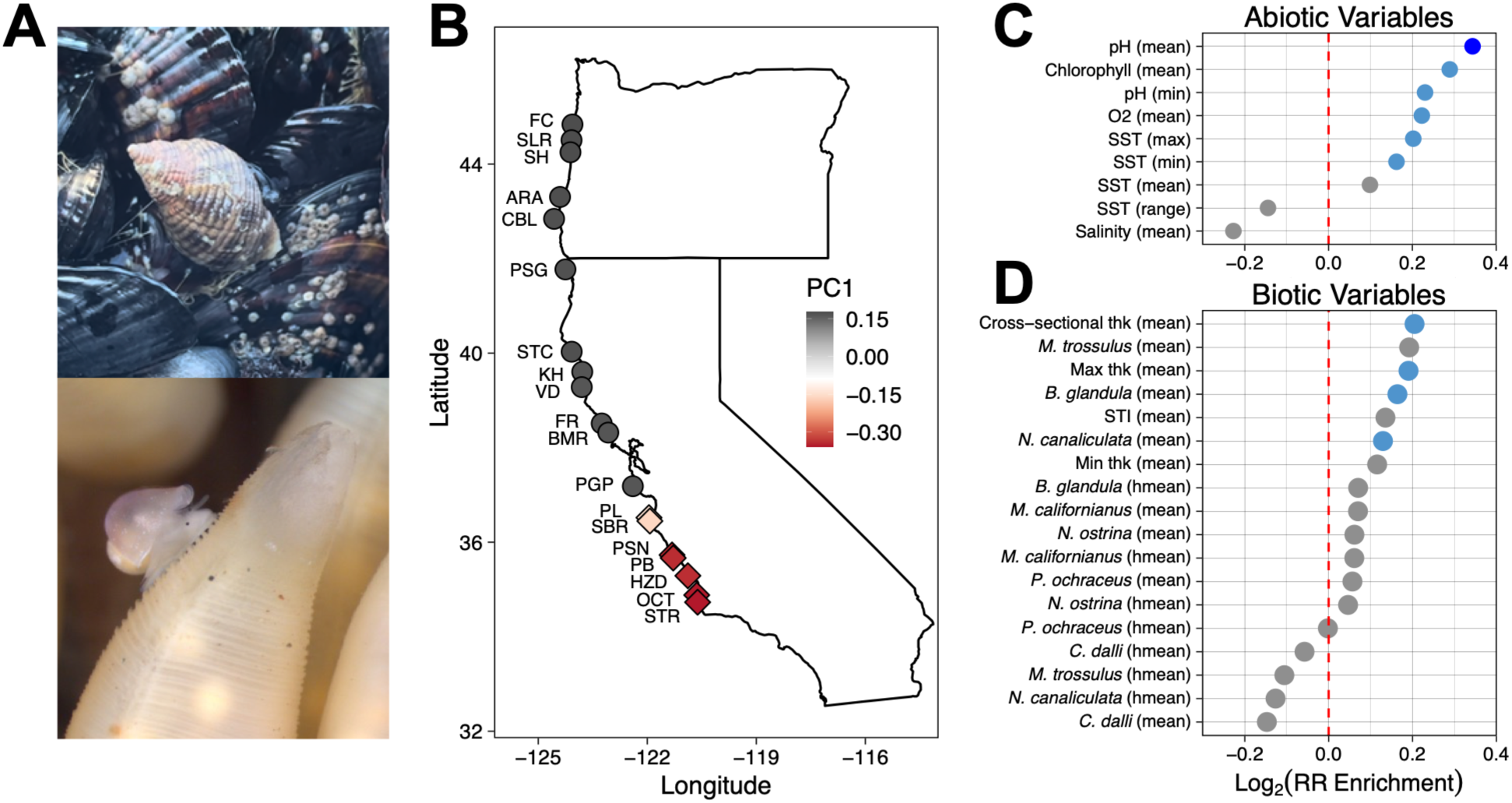
Population structure of *Nucella canaliculata* and environmental associations between *N. canaliculata* genetic variation and several abiotic and biotic variables. A) Image of an adult *N. canaliculata* in a mussel bed (top) and a newly-hatched juvenile crawling on an egg capsule (bottom). B) Population structure (PC1 of a principal component analysis) of the 19 *N. canaliculata* populations. See **Table S1** for an explanation of the site codes and coordinates. Relative rate (RR) of model enrichment for the abiotic (C) and biotic (D) ecological models. Each point represents a Generalized Linear Model (GLM) between *N. canaliculata* allele frequency and a given ecological variable with PC1 to account for demography. For each ecological variable, model enrichment is relative to 100 permutations. Models are ordered based on the strength of relative enrichment of the real signal (relative by permutation) and colored by significance – models where the mean ± 1, or 2, times the standard deviation did not include 0 are in light blue and dark blue, respectively. The summary statistic for each variable is in parentheses (hmean = harmonic mean). See **Table S2** for descriptions of the ecological variables.

Furthermore, the California Current System is an eastern boundary upwelling system with a persistent spatial mosaic of abiotic variables including temperature and pH (24, 25). These conditions are known to strongly influence the physiology and shell structure of calcifying marine invertebrates, including both *Nucella* dogwhelks (19, 26) and their prey (27–29). However, the relative influence of these abiotic forces directly on *N. canaliculata* or indirectly through the modification of prey species traits that interact with *N. canaliculata* is unclear.

Here, we integrate population genomic data with biotic and abiotic datasets to identify the ecological drivers most strongly associated with patterns of adaptive genomic variation. By combining two large-scale biogeographic datasets describing biotic interactions with environmental data on oceanic conditions, we test how spatial variation in ecological conditions shapes local adaptation across a large swath of the species range. Our ecological datasets include climatic data on oceanic conditions, biotic data on the abundance of *N. canaliculata* prey, competitors, and predators from a large-scale monitoring program on Pacific Coast rocky shores (i.e., MARINe: Multi-Agency Rocky Intertidal Network [30]), and a new dataset on the shell traits of the rocky shore foundation mussel species, *M. californianus*. Specifically, we address the primary questions: 1) What are the abiotic and biotic drivers of geographic patterns of adaptive genomic variation in a low dispersing marine species? 2) What are the genomic bases underlying patterns of local adaptation to varying selection pressures? 3) Will future environmental change erode existing patterns of local adaptation, or can populations maintain adaptation through evolutionary responses?

## Results

### Abiotic and biotic drivers of adaptive genomic variation

To identify which ecological selective forces are driving patterns of adaptive variation in *N. canaliculata*, we integrated population genomic data from 19 populations spanning 1,500 km of the west coast of North America ([15]; **Fig 1B**; **Table S1**), with large-scale datasets describing abiotic conditions and biotic interactions (**Table S2**), including a newly generated dataset of geographic variation in shell traits of the mussel prey species *Mytilus californianus* (**Table S3**). We tested associations between genome-wide allele frequency variation and 27 ecological variables (9 abiotic, **Fig 1C**; 18 biotic, **Fig 1D**), encompassing oceanographic conditions and known interacting species, while simultaneously accounting for underlying phylogeographic structure (see two major phylogeographic clusters in **Fig 1B**; [15]). The top four ecological models included abiotic variables, with the best fit model being mean pH – a variable known to be strongly linked to mollusc growth and calcification (26, 27) – followed by mean chlorophyll, minimum pH, and mean O_2_ (**Fig. 1C**). The mean pH model was statistically enriched compared to the null permutations (mean enrichment = 0.344, 95% CI: 0.320-0.368; **Fig. S1**), highlighting ocean carbonate chemistry as a major driver of local adaptation in *N. canaliculata*.

Multiple biotic models stood out as important drivers of spatial variation in *N. canaliculata* genetic diversity (**Fig. 1D**). The biotic model with the greatest relative rate of model enrichment was *M. californianus* cross-sectional thickness (mean enrichment = 0.205, 95% CI: 0.176-0.234; **Fig. S1**), a variable previously associated with variation in phenotypic traits in *N. canaliculata* (18). The biotic model with the second highest level of model enrichment was mean *M. trossulus* abundance, followed by maximum mussel shell thickness, and mean *Balanus glandula* abundance (**Fig. 1D**). These findings echo previous results from a reciprocal transplant study that indicated that *N. canaliculata* is locally adapted to the abundance of its preferred prey (17).

### Regions of the genome enriched for outlier loci associated with mean pH

To identify the genetic bases of adaptation to spatial variation in mean seawater pH (**Fig. 2A**), we conducted a genotype-environment association scan while accounting for demographic history (31, 32). Our genomic scan identified 1,828 outlier single nucleotide polymorphisms (SNPs), which were disproportionately enriched among missense and intronic variants (**Fig. S2**), suggesting an important role of both protein-coding and regulatory variation in pH adaptation.

**Figure 2.**
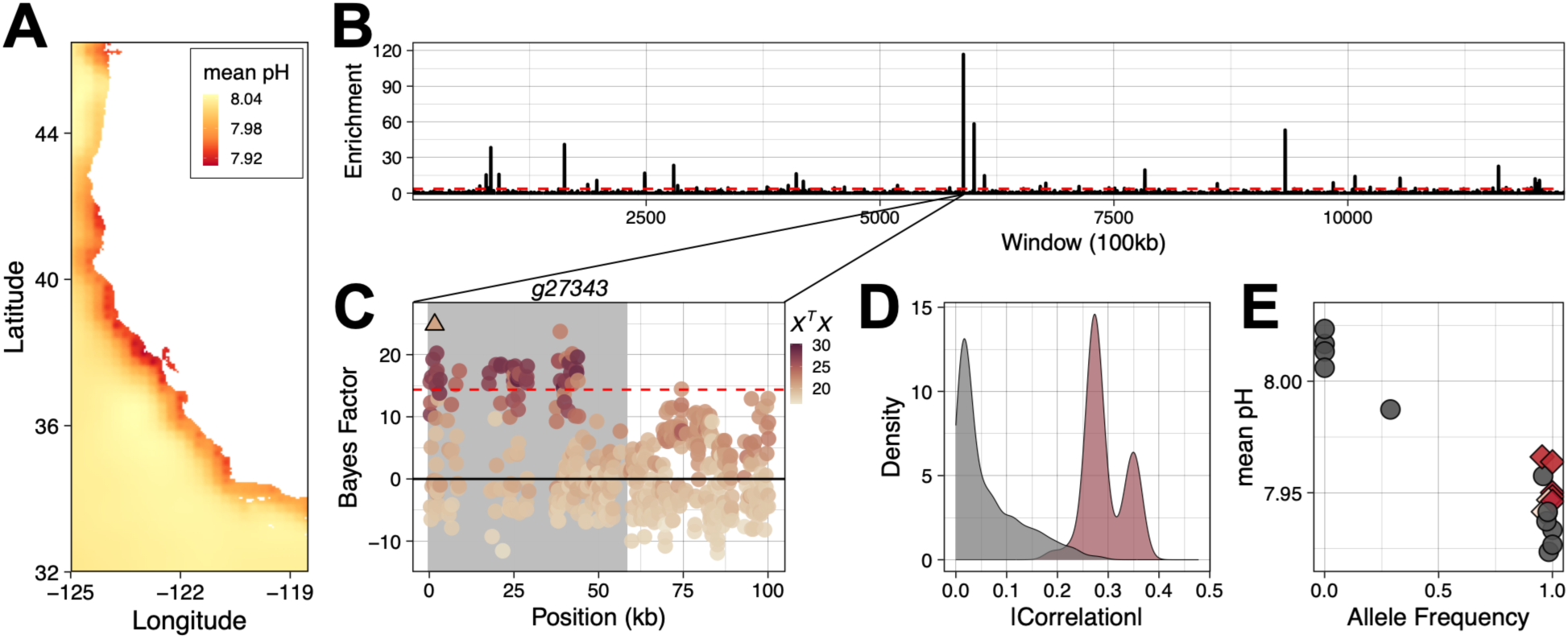
Signatures of selection associated with mean pH. A) Mean pH along the coastline from Bio-Oracle. B) Genome scan for windows enriched with outlier SNPs associated with mean pH. The x-axis is the window position along the genome and the y-axis is the -log10 transformed p-value of the enrichment test. C) Bayes Factor values and *X^T^X* for the SNPs within the top outlier window identified in B (binomial test, *P-val* = 1.99×10^-117^). The grey region indicates the position of *g27343*, which encodes for a MAM domain-containing protein and begins at the start of the scaffold. Our threshold for statistical significance (dotted red line in panels B and C) was the upper 99.9% value of the BF value from 10 simulations of pseudo-observed data (POD; BF_POD_ = 14.368). D) Absolute correlation between mean pH and the residual structure (i.e., the residuals from a model that corrected for demography and latitude) of the outlier loci in gene *g27343* that beat the POD threshold in brown (Pearson’s correlation; mean = 0.2930, sd = 0.0396) compared with a random sample of 1,000 SNPs in grey with similar mean corrected allele frequencies (mean = 0.0621, sd = 0.0657). E) Allele frequencies for the top SNP identified as a triangle in panel C. The color gradient and shapes reflect demography (PC1) as in Fig. 1B.

Furthermore, outlier SNPs were strongly clustered within 84 genomic regions rather than randomly distributed across the genome (**Fig. 2B**). Together, these patterns suggest that, while adaptation to spatial variation in pH is polygenic, it is disproportionately shaped by a small number of loci with large effects.

One window in particular stood out as being highly enriched for SNPs associated with mean pH (**Fig. 2B**). Within this window, 51 SNPs within gene *g27343* had elevated Bayes Factors (BF) and genetic differentiation (i.e., *X^T^X*) relative to the rest of the window (**Fig. 2C**). Using homology based local alignment, we identified this protein as a MAM (meprin/A5-protein/protein-tyrosine phosphatase *μ*) domain-containing protein present in the genomes of several mollusc species (**Table S4**). Proteins containing these domains are important for cell-to-cell adhesion (33) and have been linked to biomineralization processes in marine invertebrates (34, 35). Additionally, recent transcriptomic research on the sensitivity of atlantid heteropods (pelagic snails) to changing environmental conditions indicated that pH impacts the differential expression of MAM domain-containing proteins (36, 37).

The residual population structure of the 51 outlier SNPs in gene *g27343* had greater absolute correlation values with mean pH (**Fig. 2D**) than a random sample of 1,000 SNPs, indicating a strong association of these SNPs with spatial variation in ocean carbonate chemistry. When subsetting this analysis to only SNPs with the highest association values (BF>18), allele frequencies displayed strong negative correlations with mean pH (**Fig. S3**, see **Fig. 2E** for the top outlier SNP). Populations in northern California, which had the lowest mean pH values, had high allele frequencies. In contrast, populations in Oregon, which had higher mean pH values, displayed low allele frequencies or were fixed for the alternate allele. Lastly, populations in the southern genetic cluster (diamonds in **Fig. 2E** and **Fig. S3**), which have moderate pH values, had either very low or very high allele frequencies, depending on the SNP. Together, these results suggest that selection rather than demography is driving these patterns.

### Signals of selection associated with mussel cross-sectional thickness

To uncover signatures of selection to prey defenses, we analyzed patterns of genetic variation associated with the biotic variable of *M. californianus* cross-sectional thickness. Shell thickness varied biogeographically along the coastline, with thicker mussels north of San Francisco Bay (**Fig. 3A**). Despite this general trend, the population with the thickest mussels was a site in central California near Morro Bay (HZD).

**Figure 3.**
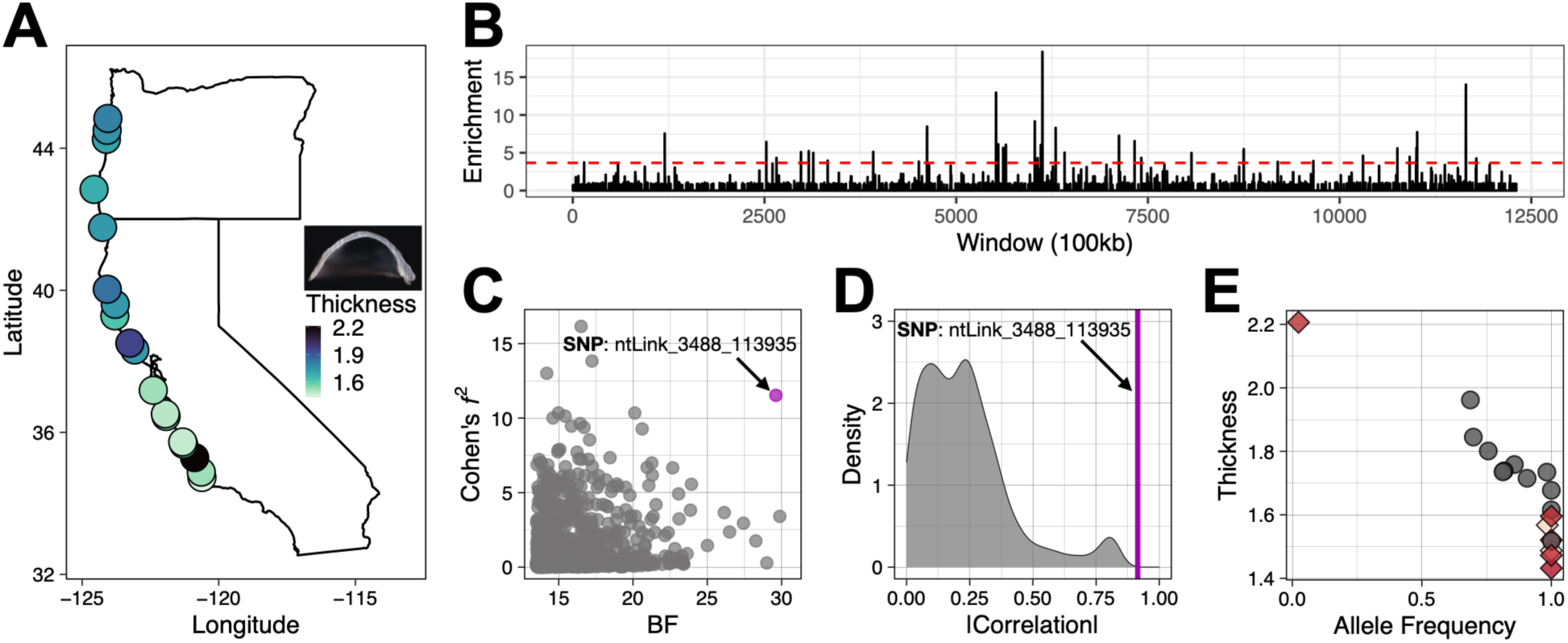
Top biotic selective driver associated with genomic variation in *Nucella canaliculata*. A) Geographic variation in *Mytilus californianus* cross-sectional thickness (mm) at ⅓ the length of the mussel along the anterior-posterior axis. Inset is an image of a mussel cross-section. B) Genome scan for windows (window size = 100kb, window step = 50kb) enriched with outlier loci associated with *M. californianus* cross-sectional thickness. The dashed red line is the 99.9% significance threshold based on the 10 simulations of pseudo-observed data (POD; BF_POD_ = 13.512). C) Association between the Bayes Factor (BF) values of the individual SNPs that beat the POD threshold and local effect size, measured as Cohen’s *f^2^*, which compared a model of demography adjusted allele frequencies with the biotic variable, demography (PC1), and latitude, and a reduced demography, and latitude-only model. D) Absolute correlation of the residual structure of the outlier SNP and *M. californianus* cross-sectional thickness (correlation = 0.9148) compared with a random sample of 1,000 SNPs (mean = 0.233, sd = 0.181). The outlier SNP (ntLink_3488_113935) is indicated in purple in panels C and D. E) Allele frequencies for the outlier SNP identified in C. The colors and shapes indicate the populations as presented in Fig. 1B.

Utilizing a genome scan that accounted for demography, we identified 52 windows that were enriched with outlier loci associated with *M. californianus* cross-sectional thickness (**Fig. 3B**). These windows appeared to be spread across the genome of *N. canaliculata* suggesting a polygenic architecture. When analyzing the effect size of individual SNPs, one SNP (ntLink_3488_113935) stood out as having both a large local effect size (Cohen’s *f^2^*) and a large BF value (**Fig. 3C**). The absolute correlation of the residual population structure of this SNP with mussel thickness was much greater than the distribution of correlations for a 1,000 SNP random sample (**Fig. 3D**). Further, the allele frequencies of this SNP were fixed for the reference allele in 6 of the sites in the southern genetic cluster (diamonds in **Fig. 3E**) but displayed clinal variation in the northern genetic cluster (circles in **Fig. 3E**). Notably, the allele frequency of the reference allele in the central California site near Morro Bay (HZD), which had the thickest mussels, was near extinction. Given the mosaic nature of this selection pressure along the coastline, these results suggest that selection is driving this pattern of genetic variation.

This outlier SNP is in an intergenic region between genes *g26430* and *g26431* – the former is an uncharacterized protein while the latter has been identified as a thiamine transporter SLC35F3 (solute carrier family 35 member f3-like protein) in several gastropod taxa (**Table S4**). The active form of thiamine (vitamin B1) is an enzyme cofactor that plays a critical role in regulating energy availability and cell metabolism (38). Thiamine deficiencies are a problem in many animals including several species of fish, crustaceans, and molluscs, with optimal levels enhancing organismal growth and feeding rates (38, 39). The process of drilling by muricid gastropods is energetically demanding (21), thus, we hypothesize that the regulation of gene *g26431* is important for modulating energy availability during this chemo-mechanical mode of feeding.

### Genetic simulations of current and future ecological conditions

To contextualize the patterns of local adaptation observed (**Fig. 2E**; **Fig. 3E**), we performed population genetic simulations of the top outlier SNP associated with both ecological stressors (40). The goal of these simulations was to emulate the strengths of selection required to recapitulate the allele frequency patterns observed using population genetic principles and simulated expectations. For both ecological stressors, we simulated a range of migration rates, population sizes, and parameters that dictate the structure of the selection curve. Based on our genetic simulations that best approximate the patterns seen in the real data, selection is over 30 times greater for mean pH (|*s*| = 0.0371 ± 0.0158; **Fig. S4**) than shell thickness (|*s*| = 0.0012 ± 0.0008; **Fig. S5**), and, consistent with organismal life history of limited dispersal, the levels of migration are low (m = 10^-4^) among populations.

Ocean acidification is forecasted to reduce pH (**Fig. 4A**), threatening the fitness of calcifying marine organisms (27) and possibly disrupting the patterns of local adaptation identified in *N. canaliculata* (**Fig. 2**). To assess the adaptive potential of *N. canaliculata* to these changing conditions, we performed population genetic simulations to model how genetic variation may change in response to declining pH. By 2032 all simulated populations were projected to have mean pH values below the selection threshold identified in the present-day model (**Fig. S6**).

**Figure 4.**
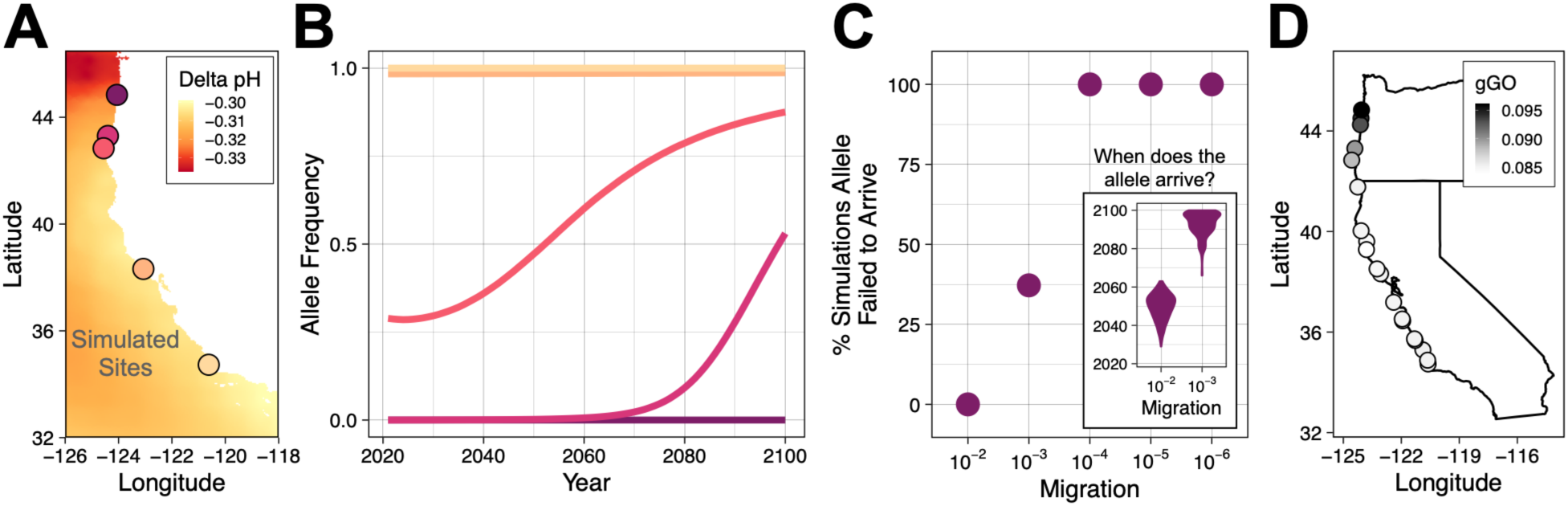
Disruption in local adaptation predicted as a result of future ocean acidification. A) Projected change in mean pH based on data from Bio-Oracle. Future values of mean pH are for the decade 2090–2100 under the SSP5-8.5 scenario. Population genetic simulations had 19 populations, but for visualization purposes, 5 simulated populations are shown. B) Estimated allele frequencies (mean of 500 iterations) for simulated dogwhelk populations for 2020–2100 in response to changing selection pressures and decreases in mean pH, assuming a migration rate of 10^-4^, a biologically realistic value for the species given their benthic development. Colors indicate the populations as displayed in A. C) Percent of simulation at which the reference allele did not reach the northernmost simulated population for a range of migration rates (N=500 per migration rate). Inset is the year at which the reference allele would first reach the northernmost population. D) Geometric genomic offset (gGO) values for the *N. canaliculata* populations based on scaled projections of mean pH for 2090–2100 and the entire Pool-Seq data.

These decreases in mean pH resulted in the reference allele, which had been locally adapted to low pH conditions (**Fig. 3E**), spreading northward (**Fig. 4B**). However, by 2100, the northernmost population did not yet have the allele, despite the simulated increased selection due to declining pH (*s* = 0.259; **Fig. 4B**). As expected, this supports that migration is critical for genetic rescue to occur. To assess these dynamics further, we performed simulations to understand the impact that a range of migration rates would have on the spread of the allele through the populations. When the migration rate was increased (m ≥ 10^-3^), the movement of this allele across space quickened such that the allele was identified in the northernmost population prior to 2100 (**Fig. 4C**). However, even at a migration rate of 0.01, the allele didn’t arrive until the middle of 2049, meaning 17 years of reduced fitness, which exceeds the relatively short lifespan of *N. canaliculata*. These results highlight the importance of considering demography when assessing the potential for an adaptive response to rapid environmental change.

Lastly, we performed genomic offset analyses to forecast genome-wide maladaptation to future ocean acidification conditions (**Fig. 4D**). All 19 populations displayed measures of future maladaptation, but the three populations in central Oregon had the greatest geometric genomic offset (gGO) values with a mean of 0.0943 (**Fig. 4D**). These results suggest that ocean acidification will disrupt current genotype-environment associations genome-wide. Notably, the pattern of genomic offset values closely mirror those of the single-locus genetic simulations, emphasizing that the strong association identified between pH and genetic variation captures biologically meaningful components of adaptation that will become maladapted with ocean acidification.

## Discussion

We found striking genomic signatures of local adaptation shaped by both abiotic and biotic selective pressures in a coastal dogwhelk, with large effect loci underlying both patterns of adaptive variation. Leveraging these genomic signals of local adaptation, we used predictive forecasting and forward simulations to show that populations differ substantially in their vulnerability to future environmental change, and that the demographic conditions maintaining current patterns of local adaptation may paradoxically constrain adaptation in the future.

Landscapes are composed of heterogeneous ecological conditions that simultaneously affect the fitness of organisms. Ecologists have long recognized that a species’ niche is a consequence of not just habitat suitability as characterized by abiotic environmental variables (41–43). Species interactions are also critical to dictating the distribution and abundance of a species, and these biotic drivers can be key selective forces that generate patterns of adaptive variation across a landscape (44–46). However, most large-scale genomic studies only consider abiotic forces as selective drivers of contemporary adaptive variation (47). Our results in *N. canaliculata* reveal parallel signatures of adaptation associated with both variation in seawater pH (**Fig. 2**) and prey shell thickness (**Fig. 3**), suggesting that populations are adaptively responding to both the physical and biological environment. The magnitude and consistency of these genomic associations indicate that local adaptation is a prominent feature of this system despite ongoing gene flow, corroborating previous phenotypic studies in this system (14–20).

Notably, the patterns of genetic differentiation that we identified do not appear to be driven by demography. The mosaic nature of upwelling in the California Current System creates spatial hotspots of exposure to low pH, rather than a latitudinal gradient in pH conditions (**Fig. 2A**; [25]). These oceanographic conditions impact the growth, survival, and calcification of marine molluscs (19, 26–29). Here, we identified that *N. canaliculata* is both directly adapted to spatial variation in pH (**Fig. 2**) and indirectly adapted to these conditions via environmentally induced changes in the traits of the mussel prey species *M. californianus* (**Fig. 3**). These results demonstrate the ecological complexities of adaptation in natural systems, and the importance of incorporating multiple dimensions of a species realized niche into studies of adaptive variation.

Our genome-wide forecasting analyses suggested that *N. canaliculata* populations are not equally positioned to cope with future environmental change. In response to ocean acidification, northern populations exhibited larger predicted genomic offsets (**Fig. 4D**), indicating a heightened risk of maladaptation under projected pH conditions. These results illustrate that contemporary adaptation may provide little information about future evolutionary resilience and highlight the importance of identifying geographic hotspots of vulnerability. However, these genomic offset approaches only estimate the amount of genetic change required to maintain current fitness-environment relationships (7, 48, 49). Thus, they do not incorporate or model the evolutionary processes that could alleviate maladaptation, including selection on standing genetic variation and gene flow (48).

Local adaptation results from the balance between selection and migration, with spatially divergent selection promoting population differentiation and gene flow homogenizing genetic variation (1–3, 50). In marine environments local adaptation was historically regarded as a rare phenomenon, since most marine invertebrates are characterized by high dispersal ability and gene flow, given the extended pelagic duration of most planktonic larvae (51). Notably, our population genetic simulations suggest that in *N. canaliculata* the observed contemporary genomic patterns are most consistent with moderate selection acting in conjunction with restricted migration among populations. Under this scenario, locally beneficial alleles are maintained because gene flow is insufficient to homogenize genetic diversity across the populations. These findings underscore the importance of demographic processes in preserving adaptive variation across the species range.

Unexpectedly, our population genetic simulations also revealed an evolutionary tension between the processes that generate local adaptation and those that facilitate adaptation to environmental change. The low migration rate that currently preserved adaptive differentiation among populations also limited the capacity of beneficial alleles to move across the linear landscape (**Fig. 4**). As environmental conditions shifted and pH declined, the northern populations quickly became maladapted to pH, causing the allele that currently occurred at high frequencies in the southern populations to become favored, suggesting that adaptation from standing genetic variation should, in principle, be possible. However, when migration was limited, the allele spread too slowly to track the pace of environmental change, resulting in a growing mismatch between population genetic composition and local selective conditions. As a result, by 2100 the northernmost population had yet to receive the beneficial allele (**Fig. 4C**). Consequently, maladaptation risk was not driven solely by the magnitude of environmental change but also by constraints on the large-scale redistribution of adaptive alleles. Rather than facilitating evolutionary rescue via maintaining higher levels of genetic diversity, as suggested in many conservation studies (52–53), local adaptation and strong population structure caused maladaptation to accumulate over time because populations were isolated from sources of beneficial alleles. Under the assumption that this locus is a bellwether for genomic rescue, these findings suggest that the northern populations are at risk of possible extirpation. Additionally, since demographic and trait variation in *Nucella* dogwhelks can impact community dynamics (22, 23), extirpation of these northern populations may have cascading eco-evolutionary consequences. Overall, this dynamic highlights the importance of considering landscape connectivity alongside selection when forecasting evolutionary responses to global change (54).

More broadly, our results emphasize that predictions of evolutionary persistence under climate change require integrating an understanding of the ecological stressors driving spatial variation in genetic diversity, with information about the demographic processes governing population connectivity. Studies frequently treat local adaptation as evidence of evolutionary potential since it maintains ecologically important genetic variation (52, 53, 55). Yet, our findings demonstrate that the same forces that generate adaptive variation may ultimately constrain an adaptive response to rapid environmental change. Identifying when adaptive alleles exist elsewhere in a species range, but cannot spread rapidly enough to reach vulnerable populations, may prove critical to conserving biodiversity under rapid global change (56, 57). Overall, our findings reveal a fundamental paradox of adaptation: the restricted gene flow that maintains local adaptation across heterogeneous landscapes can simultaneously impede the spread of beneficial standing genetic variation, causing vulnerability to climate change to increase precisely because populations have become so well adapted to their historical environments.

## Materials and methods

### SNP panel and species demography

We utilized a recently published Pool-Seq panel of genetic variation for 19 *N. canaliculata* populations along the Pacific coast of North America (15). This Pool-Seq dataset was generated from field sampling that occurred from October 2023 to July 2024, where adult dogwhelks (20 per population) were collected from 19 rocky headlands spanning ∼1,500km of coastline (**Table S1** for sampling locations). We performed stringent filtering of the published VCF file using the `poolfstat` R package (58), ultimately resulting in a dataset with 8,277,206 SNPs. We analyzed demography of the 19 populations with a principal components analysis (PCA) using the randomallele function in the `poolfstat` R package.

### Quantifying spatial variation in ecological selective forces

We used several ecologically relevant abiotic and biotic variables to test for associations between *N. canaliculata* allele frequencies and ecological conditions (**Table S2**). We obtained decadal abiotic data for the 19 focal field sites from Bio-Oracle: sea surface temperature (mean, max, min, range, 2010-2019), pH (mean, min, 2010-2018), O2 (mean, 2010-2018), salinity (mean, 2010-2019), and chlorophyll (mean, 2010-2018) (59, 60).

Recent research has indicated that *N. canaliculata* is adapted to a coastal mosaic of *M. californianus* shell thickness and displays geographic variation in predatory drilling ability (18). Thus, in 2023-2024, we collected live mussels ranging from 40-110mm in length from 18 of the focal field sites (**Table S1**). We collected mussels from the same wave-exposed intertidal habitats where the *N. canaliculata* were collected. We removed the tissue in the field and cleaned the shells with fresh water. Since mussel thickness increases with mussel length (61), we standardized the samples by binning the mussels into 10mm size bins and using a standard number of mussels from each size bin (**Table S3**). To quantify the selective landscape of mussel shell traits, we calculated four metrics from the left valves. First, we quantified a shell thickness index (STI) for the entire left valve of each mussel (62).

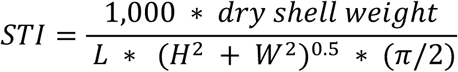

Subsequently, we cut the mussels at ⅓ their length from the anterior end, using a Dremel diamond wheel and bandsaw. This region of the shell was chosen because it is the most commonly drilled region by *N. canaliculata* (Longman & Sanford personal observation) and thus the most ecologically relevant location to quantify thickness. We scanned the mussel cross-sections and calculated three thickness metrics via imaging software (ImageJ; Java 1.8.0_172).

We quantified the shell thickness of this region as the area of the cross-section divided by the curved length of the cross-section. Additionally, we measured the maximum and minimum thickness of the cross-section, excluding the dorsal hinge for maximum thickness and the ventral lip for minimum thickness, since these areas are known to be significantly thicker and thinner, respectively. All collections were authorized under CA Fish & Wildlife Scientific Collecting Permits S-191200004-19122-001 and S-190980002-23226-001, CA State Parks permit 24-820-015 and Oregon taking permit 27976.

The second biotic dataset was species abundance data from the Multi-Agency Rocky Intertidal Network’s (MARINe) biodiversity surveys (30). We analyzed spatial variation in percent cover of potential prey species including barnacles (*Balanus glandula*, *Chthamalus dalli*) and mussels (*Mytilus trossulus*, *M. californianus*). We also considered the density of competitors (*N. ostrina*, *N. canaliculata*), and a predator (*Pisaster ochraceus*). The temporal resolution of the MARINe sites varied, plus *P. ochraceus* had a large population bottleneck as a result of sea star wasting disease in recent years (63), thus for each MARINe biotic variable we calculated the mean and harmonic mean. Sixteen of the MARINe sampling locations closely matched the locations of the *N. canaliculata* populations (**Table S1**), so genomic association analyses were performed using only these 16 of the 19 *N. canaliculata* populations.

### Generalized linear models for ecological associations

To identify associations between *N. canaliculata* allele frequencies and the abiotic and biotic ecological variables, we fit generalized linear models (GLMs) using `fastglm` [v0.0.3] (64), using a similar framework as (65). For each SNP, we calculated the mean effective coverage (*N_eff_*):

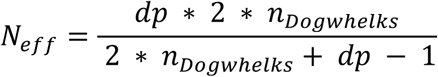

where *dp* is read depth, and *n_Do_*_gwℎ*elks*_ is the number of dogwhelks in each pool (n = 20). We fit a series of models for each SNP with *N_eff_* as the observed sample size. First, we fit a null model where allele frequencies (AF) were regressed onto their means:

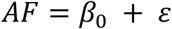

Second, since previous research indicated strong population structure (15), we fit a demographic model that regressed AF onto the first principal component (PC1).

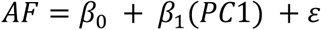

Third, we fit an ecological model that included one of the ecological variables as a covariate.

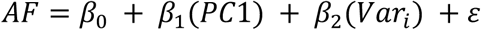

To develop null expectations for the ecological associations, we performed 100 permutations for each ecological variable. To do so, we shuffled the ecological variable across the 19 populations. These permutations preserve linkage between SNPs, thus generating null distributions of the associations. We determined the best model for each SNP using Akaike Information Criteria (AIC). To assess the relative fit of the latter two models, we performed likelihood ratio tests (LRT) between the ecological model and the demographic model.

### Model enrichment to compare ecological models

To determine the relative importance of the ecological variables, we assessed the genome-wide adaptive value of each ecological variable by comparing each ecological model to the accompanying null model generated from the permutations. For each ecological model *i*, we calculated the relative rate (*rr*) of model enrichment, across all *n* permutations for model *i*, as:

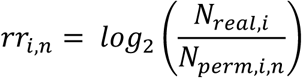

with mean and standard deviation:

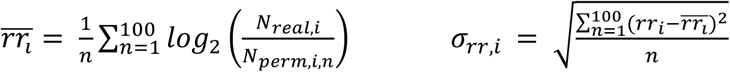

*N_real_*_,*i*_ is the number of SNPs for which a given ecological model was found to be better than the demographic model by AIC. *N_perm_*_,*i*_ is the number of SNPs for which a permuted ecological model was found to be better than the demographic model in the nth permutation.

### Outlier loci analyses

For the top abiotic and top biotic variable, we identified genomic outliers using `BayPass` [v2.41] (31). We formatted the input files for `BayPass` using the `poolfstat` R package (58), with the appropriate number of *N. canaliculata* populations based on the spatial resolution of the ecological variables. To account for the demographic history of the populations, we calculated the Ω relatedness matrix. We subsequently ran `BayPass` in the standard covariate mode with the ecological variable as a covariate. We performed 5 runs for each ecological variable, then averaged the Bayes Factors (BF) for each SNP across the iterations. To calibrate the BF term, we generated neutral distributions of the ecological associations with the genomic data by running 10 simulations of pseudo-observed data (POD). We calculated the 99.9% quantile for each POD then averaged across the simulated runs to identify a threshold for assessing significance for each ecological variable.

To identify loci associated with each ecological variable we took a two-pronged approach. First, we performed a window analysis (window size = 100 kb, step size = 50 kb, scaffolds with a minimum of 100 SNPs) on the `BayPass` datasets to identify regions within the genome that are enriched with outlier SNPs. We ranked and normalized the BF values for each SNP, such that we transformed the dataset into a uniform distribution bounded between 1 and 1/L, where L is the number of SNPs studied (66). We then summarized the windows and identified regions of the genome that contain an excess of SNPs that beat the probability threshold calculated from the PODs using binomial tests. Second, we did *post hoc* analyses that quantified the strength of the association between specific loci and the focal ecological variable. To do so, we calculated the absolute value of Pearson’s Correlation between the residual structure of the focal loci (i.e., the residuals from a linear model of the mean of the posterior distribution of the *_αij_* parameter in the 5 `BayPass` runs [a value that is closely related to the frequency of the reference allele], PC1 and latitude) and the ecological variable. We compared these correlations to those from a random sample of 1,000 loci with similar `BayPass` corrected allele frequencies. Further, we calculated the local effect size (Cohen’s *f^2^*; [67]) of specific ecological variables for individual SNPs as:

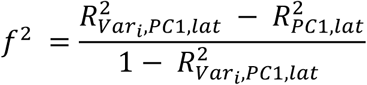

where *R*^2^ is the proportion of variance accounted for by the ecological variable (Var), demography (PC1) and latitude.

To compare the genomic architecture of adaptation to varying selective pressures, we determined if the mean pH model and the *M. californianus* cross-sectional thickness model were enriched for specific types of genetic variants using Fisher’s Exact Tests. For each model, we compared the number of each type of variant to the total number of variants for that model and the number of genome-wide variants of that type. We performed annotations of outlier loci using `SNPeff` (68). Subsequently, we did protein sequence identification using NCBI `blastp` against Molluscan taxa in the UniProtKB reference proteomes and Swiss-Prot databases.

### Population genetic simulations

To contextualize the strengths of selection needed to generate the allele frequency patterns of the top outlier SNP associated with the abiotic and biotic variables, we created single locus population genetic simulations in `SLiM` [v5.0] (40). We simulated stepping-stone populations (19 populations for mean pH and 18 for shell thickness) using a Wright-Fisher model. We explored a range of biologically realistic migration rates and population sizes for the two ecological variables, and then for comparison purposes among the models, we set migration rate and population size to fixed values for the final simulations (m=10^-4^, N = 5,000). For each model we calculated population-level selection using a logistic sigmoidal function, thus allowing the allele to be both beneficial and deleterious depending on the environment:

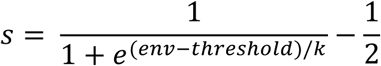

where *env* is the population-level environmental variable (mean pH or shell thickness), *t*ℎ*res*ℎ*old* is the inflection point, and *k* is the scaling factor. We did 100 simulations for each parameter combination. We incorporated pool-seq noise into the output allele frequencies after 20,000 generations using the real population-level coverage values. We performed Approximate Bayesian Computation (ABC) with `abc` [v2.2.2] (69) using the local linear regression method (tolerance = 0.1) to compare the simulated allele frequencies with the real data and identify the best-fitting parameter combination for each model. The summary statistics were mean allele frequency, Pearson’s and Spearman’s correlations between the simulated allele frequencies and the environmental variable, and Pearson’s and Spearman’s correlations between the simulated allele frequencies and the real allele frequencies. Simulations under the best-fitting parameter combinations (mean pH: *threshold* = 7.991, *k* = 0.265; shell thickness: *threshold* = 1.81, *k* = 38) closely reproduced the observed allele frequencies (**Fig. S4**; **Fig. S5**).

### Simulations of future genetic change in response to ocean acidification

To determine how the observed allele frequency pattern may change as a result of ocean acidification and projected decreases in pH, we performed future genetic simulations using `SLiM` [v5.0] (40). We used the decadal estimates of future (2020–2100) mean pH from Bio-Oracle (59, 60) under the SSP5-8.5 scenario for each population to generate population-level regression equations (**Fig. S6**). We used these equations to calculate yearly mean pH and subsequently selection per population. We started the simulations with the population allele frequencies set to the empirical values, then ran the simulation for 80 years to represent the temporal change that would occur from 2020–2100. We ran 500 simulations then averaged across the iterations. Lastly, we varied migration rate (10^-2^, 10^-3^, 10^-4^, 10^-5^, and 10^-6^) to compare the time it would take for the beneficial allele that is currently present in the southern populations to move northward and reach the northernmost simulated populations.

### Quantifying genomic offset

To calculate the genome-wide adaptive disruption anticipated from future environmental change we performed genetic offset analyses in `BayPass` (31, 70) using future projections of mean pH from Bio-Oracle (59, 60). This analysis quantified geometric genomic offset (gGO; [49]) based on scaled values of mean pH. We used the SNP regression coefficients from `BayPass`, thus accounting for the effect of current mean pH on the distribution of allele frequencies while corrected for population structure. The gGO was then calculated with respect to mean pH predicted for 2090–2100 under the SSP5-8.5 scenario for each of the 19 populations.

## Acknowledgements

Funding for this study was provided by a National Science Foundation postdoctoral fellowship (OCE-2307933) awarded to E.K. Longman. We thank the Vandenberg Space Force Base, the UC Natural Reserve System, and the Abbott family for providing access to field sites. We are grateful to N. Longman and J. Fosnight for their assistance with mussel collections, to W. Walker, S. Boutilier, M. Hilzenrath, and O. Ward for their help collecting the mussel morphology data, and to the Vermont Advanced Computing Center (VACC; https://www.uvm.edu/vacc) for providing computational resources that contributed to this publication.

## Data Sharing Plan

Code for analyzing the data can be found at https://github.com/emily-longman/Nucella_can_Seascape. The mussel shell dataset can be found on Zenodo (71).

## References

1. T. J. Kawecki, D. Ebert, Conceptual issues in local adaptation. Ecology Letters, 7, 1225– 1241 (2004).

2. F. Blanquart, O. Kaltz, S. L. Nuismer, S. Gandon, A practical guide to measuring local adaptation. Ecology Letters, 16, 1195–1205 (2013).

3. O. Savolainen, M. Lascoux, J. Merilä, Ecological genomics of local adaptation. Nature Reviews Genetics, 14, 807–820 (2013).

4. K. E. Lotterhos, The paradox of adaptive trait clines with nonclinal patterns in the underlying genes. Proceedings of the National Academy of Sciences, 120, e2220313120 (2023).

5. S. Hoban, J. L. Kelley, K. E. Lotterhos, M. F. Antolin, G. Bradburd, D. B. Lowry, M. L. Poss, L. K. Reed, A. Storfer, M. C. Whitlock, Finding the genomic basis of local adaptation: pitfalls, practical solutions, and future directions. The American Naturalist, 188, 379–397 (2016).

6. S. J. Franks, A. A. Hoffmann, Genetics of climate change adaptation. Annual Review of Genetics, 46, 185–208 (2012).

7. M. C. Fitzpatrick, S. R. Keller, Ecological genomics meets community-level modelling of biodiversity: Mapping the genomic landscape of current and future environmental adaptation. Ecology Letters, 18, 1–16 (2015).

8. J. M. Alexander, D. Z. Atwater, R. I. Colautti, A. L. Hargreaves, Effects of species interactions on the potential for evolution at species’ range limits. Philosophical Transactions of the Royal Society B: Biological Sciences, 377, 20210020 (2022).

9. L. Bernatchez, A. L. Ferchaud, C. S. Berger, C. J. Venney, A. Xuereb, Genomics for monitoring and understanding species responses to global climate change. Nature Reviews Genetics, 25, 165–183 (2024).

10. B. A. Menge, G. M. Branch. “Rocky Intertidal Communities.” in Marine Community Ecology, M. D. Bertness, S. D. Gaines M. E. Hay, Eds. (Sunderland, MA: Sinauer Associates, 2001).

11. J. H. Connell, The influence of interspecific competition and other factors on the distribution of the barnacle Chthamalus stellatus. Ecology, 42, 710–723 (1961).

12. R. T. Paine, S. A. Levin, Intertidal Landscapes: Disturbance and the Dynamics of Pattern. Ecological Monographs, 51, 145–178 (1981).

13. W. P. Sousa, Experimental investigations of disturbance and ecological succession in a rocky intertidal algal community. Ecological Monographs, 49, 227–254 (1979).

14. E. Sanford, M. S. Roth, G. C. Johns, J. P. Wares, G. N. Somero, Local selection and latitudinal variation in a marine predator-prey interactions. Science, 300,1135–1137 (2003).

15. E. K. Longman, J. C. Nunez, E. Sanford, M. H. Pespeni, Geographic divergence in population genomics and shell morphology reveal history of glacial refugia in a coastal dogwhelk. Proceedings of the Royal Society B: Biological Sciences, 293 (2026).

16. E. Sanford, D. J. Worth, Genetic differences among populations of a marine snail drive geographic variation in predation. Ecology, 90, 3108–3118 (2009).

17. E. Sanford, D. J. Worth, Local adaptation along a continuous coastline: prey recruitment drives differentiation of a predatory snail. Ecology, 91, 891–901 (2010).

18. E. K. Longman, E. Sanford, Biogeographic variation in mussel shell thickness and drilling predation on rocky shores. Oecologia, 207, 126 (2025).

19. E. S. Kuo, E. Sanford, Geographic variation in the upper thermal limits of an intertidal snail: implications for climate envelope models. Marine Ecology Progress Series, 388, 137–146, (2009).

20. I. P. Neylan, E. K. Longman, E. Sanford, J. J. Stachowicz, A. Sih, Long-term anti-predator learning and memory differ across populations and sexes in an intertidal snail. Proceedings of the Royal Society B: Biological Sciences, 291 (2024).

21. F. Rovero, R. N. Hughes, G. Chelazzi, Effects of experience on predatory behaviour of dogwhelks. Animal Behavior, 57, 1241–1249 (1999)..

22. E. L. Berlow, From canalization to contingency: historical effects in a successional rocky intertidal community. Ecological Monographs, 67, 435–460, (1997).

23. J. T. Wootton, Mechanisms of successional dynamics: consumers and the rise and fall of species dominance. Ecological Research, 17, 249–260, (2002).

24. B. Helmuth, C. D. Harley, P. M. Halpin, M. O’Donnell, G. E. Hofmann, C. A. Blanchette, Climate change and latitudinal patterns of intertidal thermal stress. Science, 298, 1015– 1017 (2002).

25. F. Chan, J. A. Barth, C. A. Blanchette, R. H. Byrne, F. Chavez, O. Cheriton, R. A. Feely, G. Friederich, B. Gaylord, T. Gouhier, S. Hacker, T. Hill, G. Hofmann, M. A. McManus, B. A. Menge, K. J. Nielsen, A. Russell, E. Sanford, J. Sevadjian, L. Washburn, Persistent spatial structuring of coastal ocean acidification in the California Current System. Scientific Reports, 7, 2526 (2017).

26. K. M. Barclay, B. Gaylord, B. M. Jellison, P. Shukla, E. Sanford, L. R. Leighton, Variation in the effects of ocean acidification on shell growth and strength in two intertidal gastropods. Marine Ecology Progress Series, 626, 109–121 (2019).

27. K. J. Kroeker, R. L. Kordas, R. N. Crim, G. G. Singh, Meta-analysis reveals negative yet variable effects of ocean acidification on marine organisms. Ecology Letters, 13, 1419– 1434 (2010).

28. K. J. Kroeker, E. Sanford, J. M. Rose, C. A. Blanchette, F. Chan, F. P Chavez, B. Gaylord, B. Helmuth, T. M. Hill, G. Hofmann, M. A. McManus, B. A. Menge, K. J. Nielsen, P. T. Raimondi, A. D. Russell, L. Washburn, Interacting environmental mosaics drive geographic variation in mussel performance and predation vulnerability. Ecology Letters, 19, 771–779 (2016).

29. J. M. Rose, C. A. Blanchette, F. Chan, T. C. Gouhier, P. T. Raimondi, E. Sanford, B. A. MengeBiogeography of ocean acidification: Differential field performance of transplanted mussels to upwelling-driven variation in carbonate chemistry. PLoS One, 15, e0234075 (2020).

30. Multi-Agency Rocky Intertidal Network (MARINe). 2023. MARINe/PISCO: Intertidal: MARINe Coastal Biodiversity Surveys: Point Contact Surveys Summarized. PISCO MN

31. M. Gautier, Genome-wide scan for adaptive divergence and association with population-specific covariates. Genetics, 201, 1555–1579 (2015).

32. L. Olazcuaga, A. Loiseau, H. Parrinello, M. Paris, A. Fraimout, C. Guedot, L. M. Diepenbrock, M. Kenis, J. Zhang, X. Chen, N. Borowiec, B. Facon, H. Vogt, D. K. Price, H. Vogel, B. Prud’homme, A. Estoup, M. Gautier, A whole-genome scan for association with invasion success in the fruit fly Drosophila suzukii using contrasts of allele frequencies corrected for population structure. Molecular Biology and Evolution, 37, 2369–2385 (2020).

33. V. B. Cismasiu, S. A. Denes, H. Reilander, H. Michel, S. E. Szedlacsek, The MAM (meprin/A5-protein/PTPmu) domain is a homophilic binding site promoting the lateral dimerization of receptor-like protein-tyrosine phosphatase μ. Journal of biological chemistry, 279, 26922–26931 (2004).

34. T. Takeuchi, L. Yamada, C. Shinzato, H. Sawada, N. Satoh, Stepwise evolution of coral biomineralization revealed with genome-wide proteomics and transcriptomics. PLoS One, 11, e0156424 (2016).

35. R. L. Flores, B. T. Livingston, The skeletal proteome of the sea star Patiria miniata and evolution of biomineralization in echinoderms. BMC Evolutionary Biology, 17, 125 (2017).

36. D. Wall-Palmer, L. Mekkes, P. Ramos-Silva, L. K. Dämmer, E. Goetze, K. Bakker, E. Duijm, K. T. Peijnenburg, The impacts of past, present and future ocean chemistry on predatory planktonic snails. Royal Society Open Science, 8, 202265, (2021).

37. P. Ramos-Silva, M. L. Odendaal, D. Wall-Palmer, L. Mekkes, K. T. Peijnenburg, Transcriptomic responses of adult versus juvenile atlantids to ocean acidification. Frontiers in Marine Science, 9, 801458 (2022).

38. J. Engelhardt, O. Frisell, H. Gustavsson, T. Hansson, R. Sjöberg, T. K. Collier, L. Balk, Severe thiamine deficiency in eastern Baltic cod (Gadus morhua). PLoS One, 15, e0227201 (2020).

39. U. Wijemanna, K. J. Lee, Dietary thiamine requirement and its effects on growth and innate immunity of Pacific white shrimp (Penaeus vannamei). Aquaculture International, 32, 2999–3016 (2024).

40. B. C. Haller, R. L. Ralph, P. W. Messer, SLiM 5: Eco-evolutionary simulations across multiple chromosomes and full genomes. Molecular Biology and Evolution, 43, msaf313 (2026).

41. C. Elton, Animal ecology. (London: Sidwick & Jackson 1927).

42. G. E. Hutchinson, Concluding remarks. Cold Spring Harbor Symposia on Quantitative Biology, 22, 415–427 (1957).

43. J. H. Brown, Macroecology. (Chicago: University of Chicago Press 1995).

44. J. N. Thompson, B. M. Cunningham, Geographic structure and dynamics of coevolutionary selection. Nature, 417, 735–738 (2002).

45. S. E. Gilman, M. C. Urban, J. Tewksbury, G. W. Gilchrist, R. D. Holt, A framework for community interactions under climate change. Trends in ecology & evolution, 25, 325– 331 (2010).

46. A. L. Hargreaves, R. M. Germain, M. Bontrager, J. Persi, A. L. Angert, Local adaptation to biotic interactions: a meta-analysis across latitudes. The American Naturalist, 195, 395–411 (2020).

47. B. Dauphin, C. Rellstab, R. O. Wüest, D. N. Karger, R. Holderegger, F. Gugerli, S. Manel, Re-thinking the environment in landscape genomics. Trends in Ecology & Evolution, 38, 261–274 (2023).

48. T. Capblancq, M. C. Fitzpatrick, R. A. Bay, M. Exposito-Alonso, S. R. Keller, Genomic prediction of (mal) adaptation across current and future climatic landscapes. Annual Review of Ecology, Evolution, and Systematics, 51, 245–269 (2020).

49. C. Gain, B. Rhoné, P. Cubry, I. Salazar, F. Forbes, Y. Vigouroux, F. Jay, O. François, A quantitative theory for genomic offset statistics. Molecular Biology and Evolution, 40, msad140 (2023).

50. J. Hereford, A quantitative survey of local adaptation and fitness trade-offs. The American Naturalist, 173(5), 579–588 (2009).

51. E. Sanford, M. W. Kelly, Local adaptation in marine invertebrates. Annual Review of Marine Science, 3, 509–535 (2011).

52. S. P. Flanagan, B. R. Forester, E. K. Latch, S. N. Aitken, S. Hoban, Guidelines for planning genomic assessment and monitoring of locally adaptive variation to inform species conservation. Evolutionary applications, 11, 1035–1052 (2018).

53. M. H. Meek, E. A. Beever, S. Barbosa, S. W. Fitzpatrick, N. K. Fletcher, C. S.Mittan-Moreau, R. N. Reid, S. C. Campbell-Staton, N. F. Green, J. J. Hellmann, Understanding local adaptation to prepare populations for climate change. Bioscience, 73, 36–47 (2023).

54. K. Schiffers, E. Bourne, S. Lavergne, W. Thuiller, J. M. Travis, Limited evolutionary rescue of locally adapted populations facing climate change. Philosophical Transactions of the Royal Society B: Biological Sciences, 368, 20120083 (2013).

55. L. M. Thompson, L. L. Thurman, C. N. Cook, E. A. Beever, C. M. Sgrò, A. Battles, C. A. Botero, J. E. Gross. K. R. Hall, A. P. Hendry, A. A. Hoffmann, C. Hoving, O. E. LeDee, C. Mengelt, A. B. Nicotra, R. A. Niver, F. Pérez-Jvostov, R. M. Quiñones, G. W. Schuurman, M. K. Schwartz, J. Szymanski, A. Whiteley, Connecting research and practice to enhance the evolutionary potential of species under climate change. Conservation Science and Practice, 5, e12855 (2023).

56. S. N. Aitken, M. C. Whitlock, Assisted gene flow to facilitate local adaptation to climate change. Annual review of ecology, evolution, and systematics, 44, 367–388 (2013).

57. S. N. Aitken, J. B. Bemmels, Time to get moving: assisted gene flow of forest trees. Evolutionary applications, 9, 271–290 (2016).

58. M. Gautier, R. Vitalis, L. Flori, A. Estoup, f-Statistic estimation and admixture graph construction with Pool-seq or allele count data using the R package poolfstat. Molecular Ecology Resources, 22, 1394–1416 (2022).

59. L. Tyberghein, H. Verbruggen, K. Pauly, C. Troupin, F. Mineur, O. De Clerck, Bio-ORACLE: a global environmental dataset for marine species distribution modelling. Global Ecology and Biogeography, 21, 272–281 (2012).

60. J. Assis, S. J. Fernández Bejarano, V. W. Salazar, L. Schepers, L. Gouvêa, E. Fragkopoulou, F. Leclercq, B. Vanhoorne, L. Tyberghein, E. A. Serrão, H. Verbruggen, O. De Clerck, Bio-ORACLE v3.0. Pushing marine data layers to the CMIP6 Earth system models of climate change research. Global Ecology and Biogeography, 33, e13813 (2024).

61. J. R. Dodd, Environmentally controlled variation in the shell structure of a pelecypod species. Journal of Paleontology, 1065–1071 (1964).

62. A. S. Freeman, J. E. Byers, Divergent induced responses to an invasive predator in marine mussel populations. Science, 313, 831–833 (2006).

63. C. M. Miner, J. L. Burnaford, R. F. Ambrose, L. Antrim, H. Bohlmann, C. A. Blanchette, J. M. Engle, S. C. Fradkin, R. Gaddam, C. D. G. Harley, B. G. Miner, S. N. Murray, J. R. Smith, S. G. Whitaker, P. T. Raimondi, Large-scale impacts of sea star wasting disease (SSWD) on intertidal sea stars and implications for recovery. PLoS ONE, 13, e0192870 (2018).

64. I.C. Marschner, Glm2: fitting generalized linear models with convergence problems. The R Journal, 3, 12 (2011).

65. J. C. Nunez, B. A. Lenhart, A. Bangerter, C. S. Murray, G. R. Mazzeo, Y. Yu, T. L. Nystrom, C. Tern, P. A. Erickson, A. O. Bergland, A cosmopolitan inversion facilitates seasonal adaptation in overwintering Drosophila. Genetics, 226, iyad207 (2024).

66. K. E. Lotterhos, D. C. Card, S. M. Schaal, L. Wang, C. Collins, B. Verity, Composite measures of selection can improve the signal-to-noise ratio in genome scans. Methods in Ecology and Evolution, 8, 717–727 (2017).

67. J. E. Cohen, Statistical power analysis for the behavior sciences. Hillsdale, NJ. (Lawrence Erlbaum Associates, Inc. 1988).

68. P. Cingolani, A. Platts, L. L. Wang, M. Coon, T. Nguyen, L. Wang, S. J. Land, X. Lu, D. M. Ruden, A program for annotating and predicting the effects of single nucleotide polymorphisms, SnpEff: SNPs in the genome of Drosophila melanogaster strain w1118; iso-2; iso-3. Fly, 6, 80–92 (2012).

69. K. Csilléry, O. François, M. G. Blum, abc: an R package for approximate Bayesian computation (ABC). Methods in Ecology and Evolution, 3, 475–479 (2012).

70. L. Camus, M. Gautier, S. Boitard, Predicting species invasiveness with genomic data: Is genomic offset related to establishment probability? Evolutionary Applications, 17, e13709 (2024).

71. E. Longman, Data from “Dataset on the shell traits of the California mussel Mytilus californianus.” Zenodo. Available at https://zenodo.org/records/21983860<u>. Deposited 17 August 2026</u>.

