## Supporting Information for "Revealing the abiotic and biotic drivers of past and future local adaptation in a coastal dogwhelk"

**Table S1**. *Nucella canaliculata* and *Mytilus californianus* sampling locations. The associated MARINe site names and codes are listed. Populations are ordered from north to south.

| Population Code | Population Name | Latitude | Longitude | *M. californianus* Sampling | MARINe Site Name (Site Code) | # Years Sampled |
|---|---|---|---|---|---|---|
| FC | Fogarty Creek, OR | 44.8378 | -124.0593 | Yes | Fogarty Creek (6) | 6 |
| SLR | Seal Rock, OR | 44.5054 | -124.0848 | Yes | Seal Rock (4001) | 3 |
| SH | Strawberry Hill, OR | 44.25 | -124.11477 | Yes | Bob Creek (7) | 5 |
| ARA | Cape Arago, OR | 43.304 | -124.40155 | No | Cape Arago (8) | 5 |
| CBL | Cape Blanco, OR | 42.841 | -124.56471 | Yes | NA | NA |
| PSG | Point Saint George, CA | 41.7712 | -124.25293 | Yes | Point Saint George (6001) | 1 |
| STC | Shelter Cove, CA | 40.0301 | -124.08091 | Yes | Shelter Cove (11) | 3 |
| KH | Kibesillah Hill, CA | 39.6046 | -123.78945 | Yes | Kibesillah Hill (12) | 6 |
| VD | Van Damme, CA | 39.2809 | -123.80357 | Yes | NA | NA |
| FR | Fort Ross, CA | 38.512 | -123.25506 | Yes | Windermere Point (83) | 1 |
| BMR | Bodega Marine Reserve, CA | 38.319 | -123.074 | Yes | Bodega (14) | 8 |
| PGP | Pigeon Point, CA | 37.1851 | -122.39758 | Yes | Pigeon Point (18) | 5 |
| PL | Point Lobos, CA | 36.5194 | -121.95367 | Yes | Point Lobos (25) | 5 |
| SBR | Soberanes Point, CA | 36.4475 | -121.92899 | Yes | Garrapata (65) | 3 |
| PSN | Point Sierra Nevada, CA | 35.7289 | -121.31867 | Yes | Point Sierra Nevada (28) | 5 |
| PB | Piedras Blancas, CA | 35.6655 | -121.28677 | Yes | Piedras Blancas (68) | 3 |
| HZD | Hazards, CA | 35.2899 | -120.88384 | Yes | Hazards (31) | 4 |
| OCT | Occulto, CA | 34.8812 | -120.63994 | Yes | NA | NA |
| STR | Stairs, CA | 34.7302 | -120.61569 | Yes | Stairs (33) | 5 |

**Table S2.** Abiotic and biotic variables used in the landscape genomic analyses to test for associations between *Nucella canaliculata* genomic variation and ecological conditions. The abbreviations refer to those used in Fig. 1.

| **Abbreviation of Ecological Variable** | **Full Ecological Variable** | **Units** | **Source** | **Summary Statistics Analyzed** |
|---|---|---|---|---|
| pH | Ocean pH at surface | Unitless | Bio-Oracle | Mean, Minimum |
| Chlorophyll | Ocean chlorophyll at surface | $mg/m^3$ | Bio-Oracle | Mean |
| O2 | Dissolved oxygen at surface | $\mu mol/m^3$ | Bio-Oracle | Mean |
| SST | Sea surface temperature | Degrees C | Bio-Oracle | Mean, Minimum, Maximum, Range |
| Salinity | Ocean salinity at surface | PSU | Bio-Oracle | Mean |
| *M. trossulus* | *Mytilus trossulus* | Percent cover | MARINe | Mean, Harmonic mean |
| *B. glandula* | *Balanus glandula* | Percent cover | MARINe | Mean, Harmonic mean |
| *C. dalli* | *Chthamalus dalli* | Percent cover | MARINe | Mean, Harmonic mean |
| *M. californianus* | *Mytilus californianus* | Percent cover | MARINe | Mean, Harmonic mean |
| *N. canaliculata* | *Nucella canaliculata* | Density | MARINe | Mean, Harmonic mean |
| *N. ostrina* | *Nucella ostrina* | Density | MARINe | Mean, Harmonic mean |
| *P. ochraceus* | *Pisaster ochraceus* | Density | MARINe | Mean, Harmonic mean |
| STI | Shell thickness index | Unitless | New dataset | Mean, Harmonic mean |
| Cross-sectional thk | Integrated thickness of the mussel cross section at ⅓ the length of the mussel calculated as the area of the section divided by the curved length | mm | New dataset | Mean, Harmonic mean |
| Max thk | Maximum thickness of the cross section at ⅓ the length of the mussel disregarding the dorsal hinge | mm | New dataset | Mean, Harmonic mean |
| Min thk | Minimum thickness of the cross section at ⅓ the length of the mussel disregarding the growing lip | mm | New dataset | Mean, Harmonic mean |

**Table S3**. Distribution of *Mytilus californianus* mussel sizes used to quantify spatial variation in shell thickness among the 19 field sites. Mussels were grouped into 10mm size bins. There were only 5 mussels that were between 70 and 80mm for Kibesillah Hill (KH), thus we included an 80.8mm mussel.

| Size Bin - Lower Threshold | Size Bin - Upper Threshold | Count |
|---|---|---|
| ≥40 | <50 | 3 |
| ≥50 | <60 | 4 |
| ≥60 | <70 | 5 |
| ≥70 | <80 | 6 |
| ≥80 | <90 | 5 |
| ≥90 | <100 | 4 |
| ≥100 | <110 | 3 |

**Table S4.** UniProt BlastP results for genes *g27343* (a MAM domain-containing protein), *g26431* (a thiamine transporter SLC35F3) and *g33776* (a kinesin domain containing-protein).

| **Gene** | **Species Name** | **UnitProt Entry** | **Percent Identity** | **E-Value** |
|---|---|---|---|---|
| *g27343* | *Batillaria attramentaria* | A0ABD0JFJ9 | 60.9% | 0.0 |
| *g27343* | *Pomacea canaliculata* | A0A2T7PKQ0 | 52.0% | 0.0 |
| *g27343* | *Mytilus coruscus* | A0A6J8BBF4 | 36.3% | 0.0 |
| *g26431* | *Patella caerulea* | A0AAN8G8G5 | 42.9% | $2.0 \times 10^{-90}$ |
| *g26431* | *Biomphalaria pfeifferi* | A0AAD8AV04 | 39.15 | $1.9\times10^{-87}$ |
| *g33776* | *Littorina saxatilis* | A0AAN9B0W0 | 44.5% | $3.2 \times 10^{-113}$ |
| *g33776* | *Batillaria attramentaria* | A0ABD0KGC3 | 52.5% | $3.3 \times 10^{-101}$ |
| *g33776* | *Pomacea canaliculata* | A0A2T7PN99 | 48.7% | $1.1 \times 10^{-96}$ |

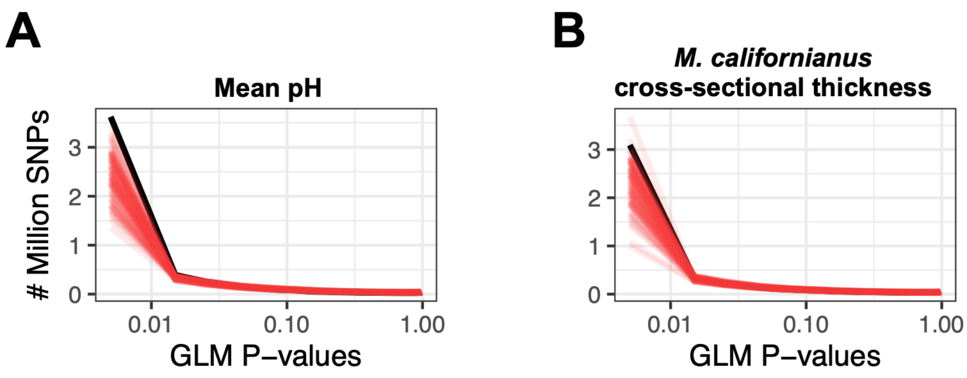


**Figure S1.** Distribution of p-values for the generalized linear model (GLM) with the real data (black line) compared to 100 permutations (red lines) for the mean pH model (A) and the *Mytilus californianus* cross-sectional shell thickness model (B).

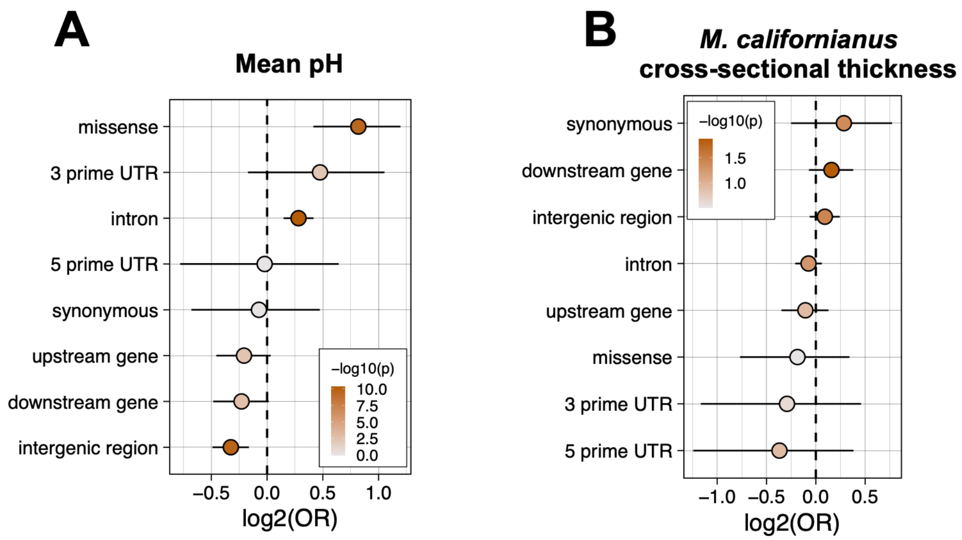


**Figure S2.** Enrichment (odds ratio [OR]) and 95% confidence intervals of the mean pH (A) and cross-sectional mussel shell thickness model (B) compared to the rest of the genome for different classes of genetic variants. Points are colored based on the -log10(*P*-value). The vertical dashed line represents the null expectation of no enrichment.

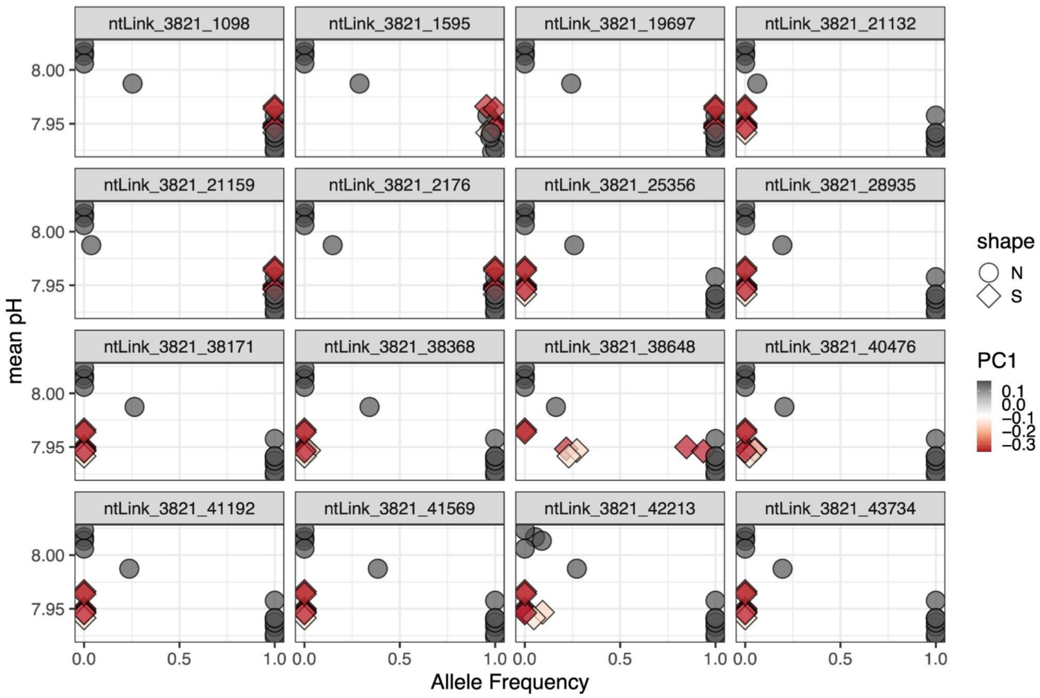


**Figure S3.** Allele frequencies of the 16 SNPs with Bayes Factors greater than 18 in gene *g27343*. Color indicates PC1 of a principal component analysis and shape differentiates the two demographic clusters - circles (north of Monterey Bay) and diamonds (south of Monterey Bay).

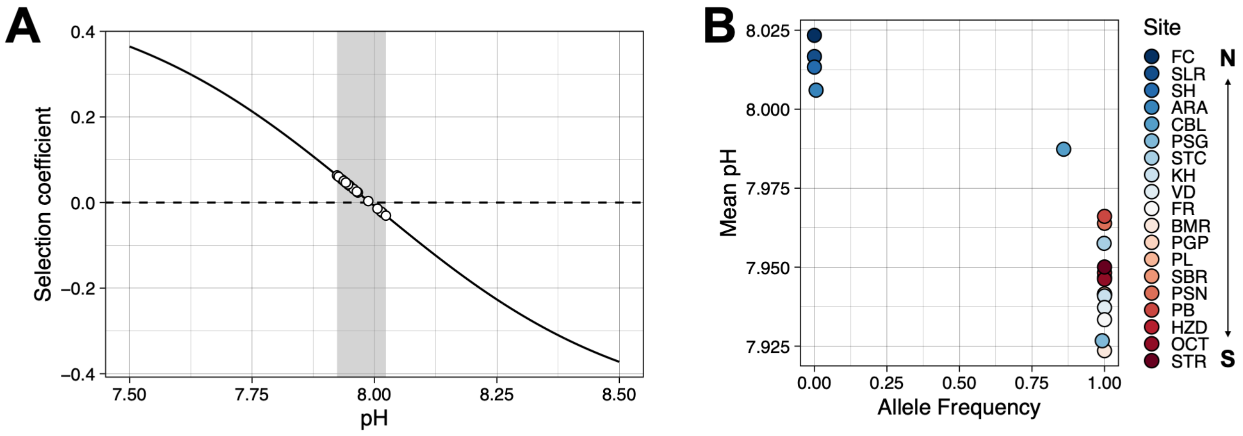


**Figure S4.** Population genetic simulation results for the best-fitting parameter combination for mean pH (*threshold* = 7.991, *k* = 0.265). A) Selection coefficients for a range of pH values that are possible in the California Current System. The grey region represents the mean pH values for the 19 *N. canaliculata* populations, with the points being the selection coefficients of the simulated populations. B) Simulated allele frequencies (mean of 100 iterations) for the best-fitting parameter combination for the 19 populations. See **Table S1** for an explanation of the site codes and coordinates.

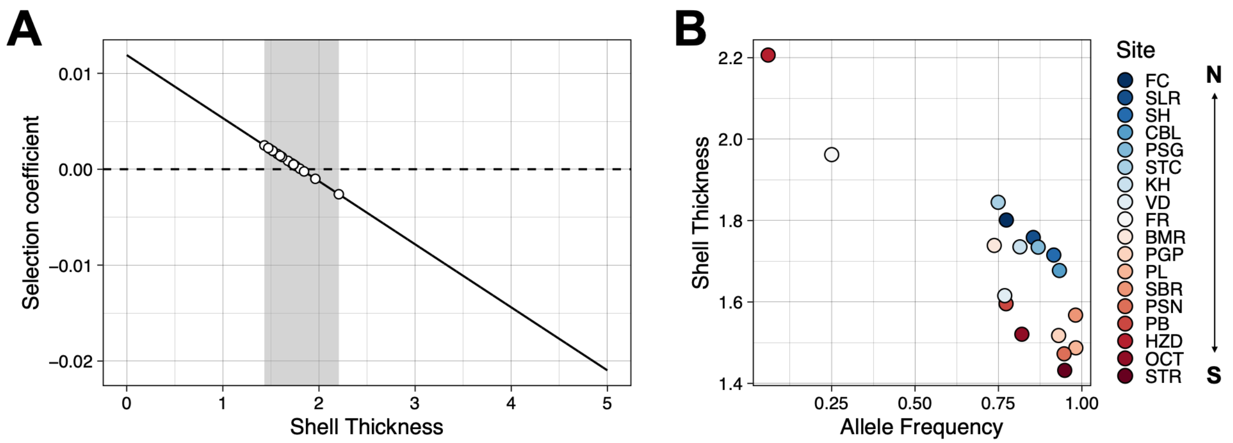


**Figure S5.** Population genetic simulation results for the best-fitting parameter combination for mussel cross-sectional shell thickness (*threshold* = 1.81, *k* = 38). A) Selection coefficients for a range of mussel shell thicknesses (mm). The grey region represents the cross-sectional shell thicknesses for the 18 *N. canaliculata* populations, with the points being the selection coefficients of the simulated populations. B) Simulated allele frequencies (mean of 100 iterations) for the best-fitting parameter combination for the 18 populations. See **Table S1** for an explanation of the site codes and coordinates.

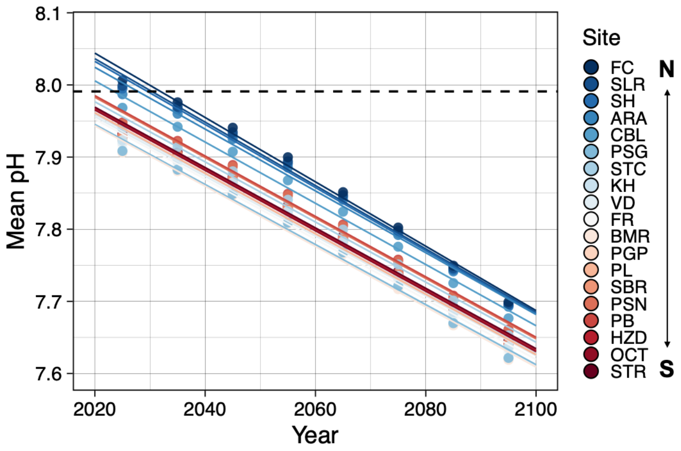


**Figure S6.** Projected mean pH values for the 19 *Nucella canaliculata* populations from Bio-Oracle under the SSP5-8.5 scenario. The x-axis starts at 2020 and displays 80 years into the future until 2100. Colored lines are linear regressions for each population. The dotted black line is the statistical threshold for selection determined in the population genetic simulations. Site abbreviations are defined in **Table S1**.
